# Fat food exacerbates post-prandial hypothalamic inflammation involving GFAP+ cells and microglia

**DOI:** 10.1101/835967

**Authors:** C. Cansell, K. Stobbe, O. Le Thuc, CA. Mosser, S. Ben-Fradj, J. Leredde, C. Lebeaupin, D. Debayle, L. Fleuriot, F. Brau, N. Devaux, A. Benani, E. Audinat, N. Blondeau, JL. Nahon, C. Rovère

## Abstract

In humans, obesity was associated with brain inflammation and glial cell proliferation. Studies in rodents showed that glial cell proliferation occurs within 24 hours of high-fat diet (HFD) consumption, before obesity development. This proliferation was mainly observed in the hypothalamus (HT), a crucial brain structure for controlling body weight. Therefore, we sought to characterize the post-prandial HT inflammatory response to 1-3-6 hours exposure to a standard diet and HFD. HFD exposure increased gene expression of astrocyte and microglial marker (GFAP and Iba1) compare to standard treated mice and induced morphological modifications of microglial cells in HT. This remodeling was associated with higher expression of inflammatory genes and differential activation of hypothalamic neuropeptides involved in energy balance regulation. DREADD and PLX5622 technologies, used to modulate GFAP-positive or microglial cells activity respectively, showed that both glial cell types are involved in hypothalamic post-prandial inflammation, but in a different time frame and with a diet specificity Thus, an exacerbated post-prandial inflammation in brain might predispose individuals to obesity and needs to be characterized to address this worldwide crisis.

## 1. Introduction

Obesity is a major public health problem that has been highlighted by the World Health Organization. In 2016, almost 40% of the worldwide population was overweight and 13% was obese (http://www.who.int/en/). The main driving force of this body weight disturbance is an energy balance dysregulation, due to kilocalorie consumption overriding energy expenditure (Gao and Horvath, 2008). Therefore, chronic exposure to the Western diet rich in saturated fat (commonly known as the high-fat diet (HFD)) is now recognized as a significant contributor to the development of obesity and its associated comorbidities, such as insulin resistance, type 2 diabetes mellitus and other chronic illnesses such as cardiovascular disease or cancer (Hotamisligil, 2006).

Excessive intake of dietary fat induces chronic low-grade inflammation in many metabolic organs including the pancreas, adipose tissue, muscle and liver, as well as in the brain (Lumeng and Saltiel, 2011). HFD consumption increases the expression of inflammatory mediators (De Souza et al., 2005; Thaler and Schwartz, 2010), especially the pro-inflammatory cytokines Interleukin-1 beta (IL-1β), Tumor Necrosis Factor alpha (TNF-α) and Interleukin 6 (IL-6), and disrupts insulin and leptin signaling in murine and primate hypothalamus (HT), a brain area involved in energy balance regulation (Arruda et al., 2011; Grayson et al., 2010; Milanski et al., 2009; Zhang et al., 2008). HFD-induced inflammatory modifications of brain signals also converge to alter levels of hypothalamic neuropeptides, suggesting a new concept where hypothalamic neuropeptides act as a link between central inflammation, dysregulation of feeding behavior and energy homeostasis, and indicating that these peptides could be potential therapeutic targets for the treatment of obesity (Timper and Bruning, 2017).

Since 2012, several studies have shown that HFD consumption induces astrogliosis and microgliosis in the arcuate nucleus (ARC) of the HT (Andre et al., 2017; Balland and Cowley, 2017; Baufeld et al., 2016; Gao et al., 2014; Guillemot-Legris et al., 2016; Kim et al., 2019; Thaler et al., 2012; Valdearcos et al., 2017; Valdearcos et al., 2014). The microglia activation that is induced by lipids and connects dietary fat and brain inflammatory signaling is begging to be seen as the missing link between the hypothalamic response to HFD and susceptibility to obesity (Andre et al., 2017; Kim et al., 2019; Valdearcos et al., 2017; Valdearcos et al., 2014). Nevertheless, it has recently been shown that HFD-induced inflammation in astrocytes may have a protective effect against the development of obesity (Buckman et al., 2015). Interestingly, hypothalamic inflammation has been described very early in response to HFD, before the onset of obesity (Balland and Cowley, 2017; Buckman et al., 2015; Kim et al., 2019; Thaler et al., 2012; Waise et al., 2015). Exposure to HFD for a 24h period increases gene expression of inflammatory astrocyte and microglia markers as well as astrogliosis in the HT (Buckman et al., 2015; Dalvi et al., 2017; Thaler et al., 2012). Microgliosis has also been found in the ARC after a 72-hour exposure to HFD (Thaler et al., 2012). These findings tempt us and others to speculate that hypothalamic inflammation may not be limited to chronic exposure to dietary fat, but may occur after consumption of a single meal (Cani et al., 2007; Emerson et al., 2017; Ghanim et al., 2009; Herieka and Erridge, 2014; Kelly et al., 2012; Khor et al., 2014). Therefore, we investigated cellular and molecular hypothalamic remodeling in mice that were exposed for the first time to a HFD for short periods of time (1h, 3h or 6h) and we compared it to mice exposed to standard diet (SD).

## 2. Materials and methods

### 2.1. Animals

8-week-old C57Bl/6J male mice (20-25 g, Janvier Labs, France) and 9-10 week-old CX3CR1eGFP/eGFP male mice (Jung et al., 2000) were housed in a room maintained at 22+/-1°C with a 12h light/12h reversed dark cycle and were acclimatized for 2-3 weeks before experiments were performed. Animals had access to water and standard diet *ad libitum* (SD; 3395 kcal/kg with 25.2% from proteins, 13.5% fat and 61.3% from carbohydrates; Safe #A03). All protocols were carried out in accordance with French standard ethical guidelines for laboratory animals and with approval of the Animal Care Committee (Nice-French Riviera, registered number 04042.02; University Paris Descartes registered number CEEA34.EA.027.11 and CEEA16-032; APAFIS#14072-2018030213588970 v6). The heterozygous CX3CR1eGFP/+ mice used for electrophysiology were obtained by crossing CX3CR1eGFP/eGFP (Jung et al., 2000) with C57BL/6J (Janvier Labs, France) wild-type mice.

### 2.2. Short-term high-fat feeding studies

Mice were food-deprived for 2h prior the onset of the dark cycle to synchronize groups. At the beginning of the dark cycle (T=0h) mice were fed either standard diet (SD; 3395 kcal/kg with 25.2% from proteins, 13.5% fat and 61.3% from carbohydrates; Safe #A03) or high-fat diet (HFD; 4494.5 kcal/kg with 16.1% from proteins, 40.9% from fat and 43% from carbohydrates; Safe #U8954P V0100). One hour, 3h, and 6h after food exposure, food intake was measured and brain and blood samples were collected. Microglia were depleted by administering the Colony Stimulating Factor 1 Receptor (CSF1R) inhibitor PLX5622 (Plexxikon, USA), formulated in AIN76A (3827.7 kcal/kg with 18.2% from proteins, 12.6% fat and 69.2% from carbohydrates; Research Diet, USA) at a dose of 1.2 g/kg for 2 weeks. Clozapine N-oxide (CNO; Sigma, France) was prepared in water at 1 mg/kg.

### 2.3. Stereotactic virus injections

Mice were anesthetized by intraperitoneal (i.p.) injection of a ketamine-xylazine mix (80 mg/kg - 12 mg/kg). They were then placed on a stereotaxic frame. AAV8/GFAP-HA-hM4G(Gi)-mCitrine (Translational Vector Core, France) virus was injected bilaterally into the medio basal HT (MBH). Stereotactic coordinates relative to bregma were: x: ±0.3 mm; y: −1.5 mm; z: +6 mm. Injections were applied at a rate of 0.5 μL/min for 1 min per side. At the end of the surgical procedures, mice received 1 mg/kg i.p. atipamezole and 5 mg/kg subcutaneous (s.c.) ketoprofen.

### 2.4. RNA isolation and quantitative PCR

Total RNA from HT was isolated using Fast Prep apparatus (Q-Biogene, France) as previously described (Le Thuc et al., 2016). First-strand cDNAs were synthesized from 2 μg of total RNA with 200 U of SuperScript III reverse transcriptase (SuperScriptIII, Invitrogen, France) in the appropriate buffer in the presence of 25 ng/μL oligo-dT primers, 0.5 mM desoxyribonucleotide triphosphate mix, 5 mM dithiothreitol, 40 U RNAsin (Promega, France). The reaction was incubated 5 min at 25 °C, then 50 min at 50 °C then inactivated for 15 min at 70 °C. Real-time PCR was performed for amplification of mouse IL-1β, IL-6, TNF-α, CCL2, CCL5 (Chemokine (C-C motif) ligand 2-5), Iba1 (Ionized calcium Binding Adaptor molecule 1), GFAP (Glial Fibrillary Acidic Protein), MCH (Melanin-Concentrating Hormone), ORX (OReXine), POMC (Pro-OpioMelanoCortin), CART (Cocaine- and Amphetamine-Regulated Transcript), NPY (Neuropeptide Y), AgRP (Agouti-Related Peptide) and GAPDH (GlycerAldehyde Phosphate DesHydrogenase) mRNA. GAPDH was used as housekeeping gene for normalization. Primers (detailed in Supplementary data, Supplementary Table 1) were purchased from Eurogentec (France).

### 2.5. Cytokine and chemokine quantification

A V-Plex multiplex assay (Meso Scale Discovery, USA) and a mouse CCL5 ELISA Ready-SET-Go (eBiosciences, France) were used to measure the levels of inflammatory mediators in mice serum according to the manufacturer’s protocol.

### 2.6. Immunohistochemical analysis

Brains were harvested from mice perfused with 4% paraformaldehyde in phosphate buffer saline (PBS) and post-fixed in the same fixative overnight at 4°C. Brain coronal sections (30 μm) were cut on a vibratome, blocked for 1h with 3% normal goat serum in PBS containing 0.1% Triton X-100 and incubated with primary antibodies overnight at 4°C. Primary anti-rabbit antibodies were against Iba1 (1:500, CP290A, B, Biocare Medical, USA) and GFAP (1:300, Z0334, Dako, Danemark). Adequate Alexa Fluor ® 488conjugated secondary antibodies were used for immunofluorescence microscopy. Sections at −1.70 mm relative to Bregma were mounted in Vectashield solution (H-1000, Vector Laboratories). 3D mosaics of 1024×1024 images were acquired with a TCS SP5 laser-scanning confocal microscope (Leica Microsystems, Nanterre, France) through a 40X/1.4 Oil immersion objective for GFAP staining and 63X/1.4 Oil immersion objective for Iba1 staining, with a z-step of 2μm. GFAP, Iba1 and soma size measurements were done on ImageJ (Schneider et al., 2012) on maximal intensity projections of these z-stacks of images. After the z-projection, a Region Of Interest (ROI) corresponding to the ARC of the hypothalamus was manually drawn on these images. GFAP and Iba1 staining areas were selected and measured (in μm^2^) in these ROIs by intensity thresholding and measurement above this threshold. The same process was applied to measure the soma size of microglial cells.

### 2.7. Acute HT slices preparation for microglia recordings

Male mice aged between 9 and 10 weeks were anesthetized with isoflurane before the brain was harvested and coronal slices through the HT were cut using a Leica VT1200 vibratome in an oxygenated (5% CO_2_ and 95% O_2_) ice-cold protective extracellular solution containing (in mM): 93 NMDG, 2.5 KCl, 1.2 NaH_2_PO_4_, 30 NaHCO3, 20 HEPES, 2 thiourea, 25 D-glucose, 5 sodium ascorbate, 3 sodium pyruvate, 10 MgCl_2_, and 0.5 CaCl_2_ (pH 7.3, 320 mOsm). Slices were then incubated in the same protective extracellular solution for 7 min at 34°C and then incubated for 30 min in artificial cerebro-spinal fluid (aCSF; pH 7.3, 310 mOsm) at 34°C containing (in mM): 135 NaCl, 2.5 KCl, 26 NaHCO_3_, 1.25 NaH_2_PO_4_, 2.5 D-glucose, 1 sodium pyruvate, 1 MgCl_2_, and 2 CaCl_2_. The slices were then maintained at room temperature (RT, 22°C) for 0.5–5 h in the regular oxygenated aCSF. After incubation, individual slices selected for best visibility of the ARC were transferred to a recording chamber on the stage of an Olympus microscope (BX50WI) with a 40x water immersion objective, equipped with a CCD camera (Hamamatsu ORCA2-AG, France). Slices were constantly perfused at RT with oxygenated aCSF (5 mL/min).

### 2.8. Electrophysiological recordings

Visually-identified eGFP-expressing microglial cells, located at least 50 μm below the slice surface, were patched in whole-cell configuration in the ARC. Micropipettes (6 to 7 MV) were filled with a solution containing (in mM): 132 K-gluconate, 11 HEPES, 0.1 EGTA, 4 MgCl_2_ (pH 7.35, osmolarity 300 mOsm). All potential values given in the text were corrected for a junction potential of 10 mV. Voltage-clamp recordings were performed using an Axopatch 200B (Molecular Devices, USA). Series resistance (Rs) as well as the cell membrane resistance (Rm), capacitance (Cm) were determined from the current response to a 10 ms depolarizing pulse of 10 mV that was performed at the beginning of each recording. These currents were filtered at 10 kHz and collected using PClamp 10 (Molecular Devices, USA) at a frequency of 100 kHz. To measure the I/V relationship, we used hyperpolarizing and depolarizing steps (from –150 to +40 mV for 50 ms), and currents were low-pass filtered at 5 kHz and collected at a frequency of 20 kHz. Currents were analyzed off line using Clampfit 10.7 (Molecular Devices, USA). To measure capacitance, currents were first fitted with a double exponential function in order to determine the time constant tau (τweighted) defined as τweighted = (A1*τ1+A2*τ2)/(A1+A2) and then capacitance was calculated as Cm=(τweighted(Rm+Rs))/((Rm*Rs)).

### 2.9. LPS mass quantitation by LC-MSMS

Lipopolysaccharide (LPS) mass concentration was determined by direct quantitation of 3-hydroxytetradecanoic acid (or 3 hM) by LC/MS/MS as previously described by Pais de Barros et al. (Pais de Barros et al., 2015). Quantitative LPS analysis using HPLC/MS/MS was performed as well as the lysate assay of its combination with the limulus amebocyte according to Pais de Barros et al. with modifications. Plasma (ca 50 μL) was mixed with saline (up to 100 μL), 4 pmol of internal standard (3-hydroxytridecanoic acid, 1 pmol/μL in ethanol) and finally hydrolyzed with 300 μL of 8 M HCl for 3h at 90°C. Total fatty acids were extracted with 600 μL distilled water and 5 mL ethyl acetate/hexane (3:2 v/v). After recovery and vacuum evaporation of the organic upper phase, fatty acids were dissolved in 200 μL ethanol and transferred into 200 μL micro-inserts for LC vials. After evaporation of ethanol under vacuum, samples were finally dissolved in 50 μL ethanol for LC/MS/MS analysis (injection volume 2 μL). Fatty acid separation was performed on an Infinity 1260 HPLC binary system (Agilent Technologies, France) equipped with a ZORBAX SB-C18 C18 50 × 2.1 mm 1.8 μm (Agilent Technologies, France) maintained at 45°C with a binary set of eluents (eluent A: 1 M ammonium formate/formic acid/water 5/1/996 v/v/v and eluent B: 1 M ammonium formate/formic acid/water/acetonitrile 5/1/44/950 v/v/v) at a flow rate of 0.4 mL/min. An 8 min eluent gradient was established as follows: from 0 to 0.5 min, the mobile phase composition was maintained at 45% B; then the proportion of B was increased linearly up to 100% over 2.5 min and maintained for 5 additional minutes. The column was equilibrated with 45% B for 5 min after each sample injection. MS/MS analysis was performed on a 6490 triple quadruple mass spectrometer (Agilent Technologies, France) equipped with a JetStream-Ion funnel ESI source in the negative mode (gas temperature 290°C, gas flow 19 L/min, nebulizer 20 psi, sheath gas temperature 175°C, sheath gas flow 12 L/min, capillary 2000 V, V charging 2000 V). Nitrogen was used as the collision gas. The mass spectrometer was set up in the selected reaction monitoring (SRM) mode for the quantification of selected ions as follows: for 3-hydroxytetradecanoic acid, precursor ion 243.2 Da, product ion 59 Da, fragmentor 380 V, collision cell 12 V, cell acceleration 2 V; for 3-hydroxytridecanoic acid, precursor ion 229.2Da, product ion 59 Da, fragmentor 380 V, collision cell 12 V, cell acceleration 2 V.

### 2.10. Triglyceride analysis

The detailed protocol is provided in the Supplementary data, Supplementary Materials and Methods. Briefly, lipids from mouse serum were extracted according to a modified Bligh and Dyer protocol with a mixture of methanol/chloroform (2:1). The lipid extract was then separated on a C18 column in an appropriate gradient. Mass spectrometry data were acquired with a Q-exactive mass spectrometer (ThermoFisher, France) operating in data dependent MS/MS mode (dd-MS2). Finally, lipids were identified using LipidSearch software v4.1.16 (ThermoFisher, France) in product search mode.

### 2.11. Statistical analysis

Displayed values are means ± SD. To test if the data set was well-modeled, a Kolmogorov-Smirnov normality test was conducted (with Dallal-Wilkinson-Lillie for P value). The ROUT method (robust regression and outlier removal) was used to identify outliers with a Q coefficient equal to 1%. Variance equality was tested using a F-test. If samples fulfilled normal distribution and variance equality criteria, comparisons between groups were carried out using an unpaired t test for single comparison and a two-way ANOVA for multiple comparisons and interaction. If samples did not follow a normal distribution or had different variances, comparisons between groups were carried out using a non-parametric Mann-Whitney U test for single comparison and non-parametric Kruskal-Wallis test with Dunn’s correction for multiple comparisons. When appropriate, comparison with the theoretical mean control (1 or 100%) was done using a non-parametric Wilcoxon signed rank test. A p-value of 0.05 was considered statistically significant. All tests were performed using GraphPad Prism 7.02 and Microsoft Office Excel. Numbers of animals and cells are given in the legends.

## 3. Results

### 3.1. A single high-fat meal increases serum levels of saturated triglycerides and endotoxins, but not those of cytokines and chemokines

Obesity and metabolic syndromes are concomitant with a state of chronic systemic inflammation (Lumeng and Saltiel, 2011) and the degree to which triglycerides (TG) increase after HFD consumption correlates with the probability of developing metabolic syndrome (Herieka and Erridge, 2014; Nogaroto et al., 2015). Because the main difference between SD and HFD is lipid content, we measured TG serum levels after short exposure to the diet. By mass spectrometry, we found that HFD consumption did not change the total TG serum levels (Fig. 1A) but led to increase the percentage of TGs containing only saturated fatty acids (Fig. 1B) compared to SD-fed mice. Previous studies have linked HFD consumption to post-prandial systemic inflammation (Cani et al., 2007; Emerson et al., 2017; Ghanim et al., 2009; Herieka and Erridge, 2014; Kelly et al., 2012; Khor et al., 2014). Therefore, we measured the serum levels of the pro-inflammatory cytokines IL-1β, IL-6 and TNFα, pro-inflammatory chemokines CCL2 and CCL5 (Chemokine (C-C motif) Ligand 2-5), as well as endotoxin (lipopolysaccharide, 3OH C14:0) after SD and HFD short-term exposure (1h, 3h and 6h). Interestingly, HFD consumption did not increase pro-inflammatory cytokine or chemokine serum levels compared to SD-fed mice over the 3 time points (Fig. 1C-G). However, 3h and 6h of HFD consumption increased endotoxin serum levels compared to SD-fed mice (Fig. 1H). As anticipated (Buckman et al., 2015), an increase in kilocalories consumed was noticeable in mice fed for 6h with HFD compared to SD-fed mice (Supplementary Fig. 1A), while the total quantity in grams of ingested food was similar between groups (Supplementary Fig. 1B). Altogether, these observations prove that short-term HFD exposure does not affect systemic levels of inflammatory markers, but increases endotoxin concentrations and the proportion of saturated TGs in the serum compared to SD-fed mice.

**Figure 1:**
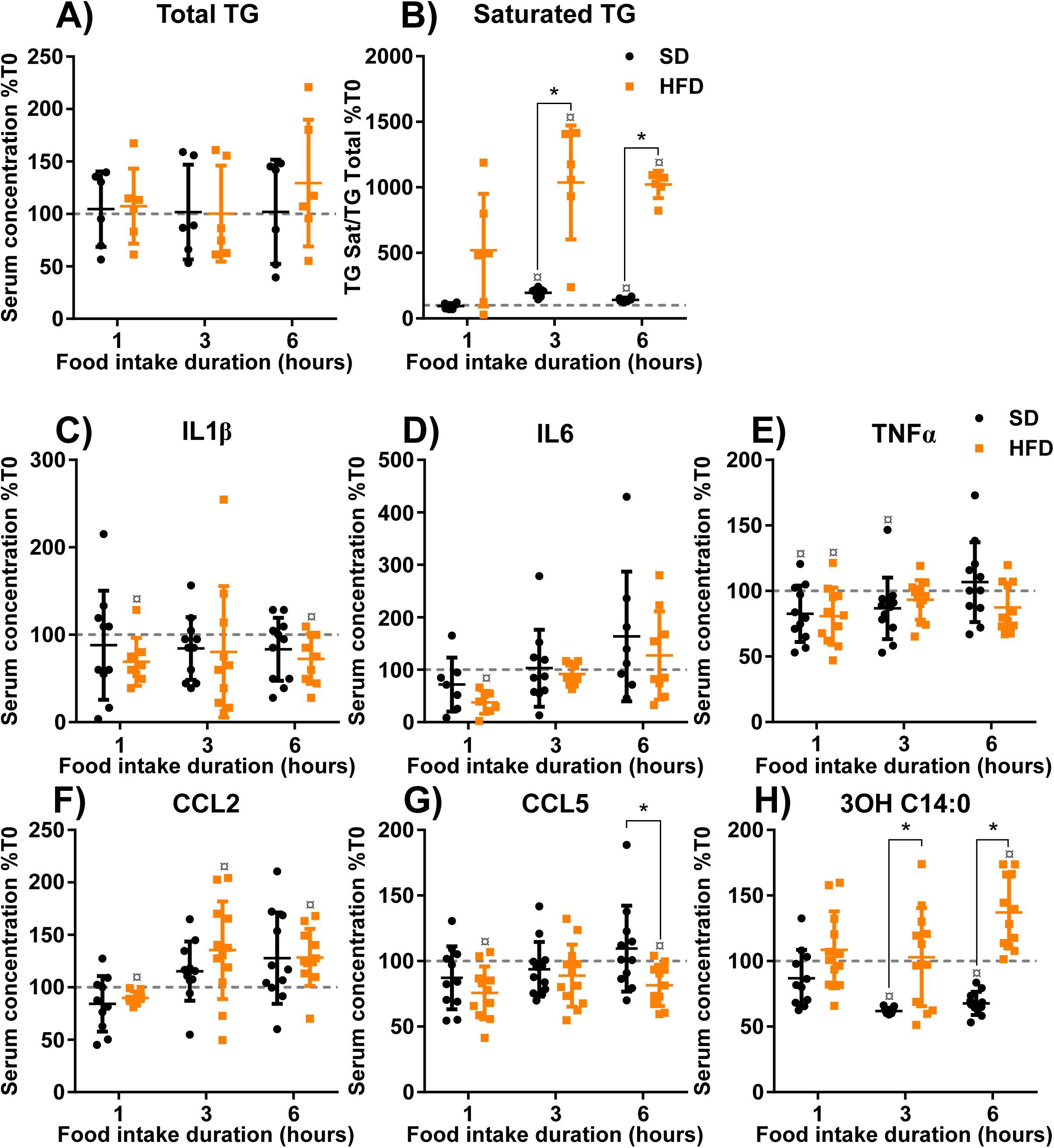
Short term high-fat diet exposure increases saturated triglyceride and endotoxin levels without affecting cytokine and chemokine levels in serum. Serum levels of total triglycerides (TG) (A), saturated TG (B), proinflammatory cytokines IL-1β (C) IL-6 (D) TNFα (E), chemokines CCL2 (F) CCL5 (G), endotoxin (3OH C14:00; H) of 8-week-old C57Bl/6J male mice fed for 1h, 3h and 6h with either standard diet (SD) or high-fat diet (HFD). Results are expressed in % compared to baseline prior to diet exposure (T0, dashed line = initial level). *p < 0.05 Mann–Whitney test or unpaired t-test SD *vs* HFD; ¤p ≤ 0.05 Wilcoxon Signed Rank test SD/HFD ≠ 100; Mean+/- SD. N=6-12.

### 3.2. A single high-fat meal induces an exacerbated inflammatory-like gene response in the hypothalamus

Given the lack of a peripheral inflammatory response in HFD-fed mice, we subsequently investigated the impact of a HFD on hypothalamic inflammatory markers. Consistent with previous studies (Andre et al., 2017; Baufeld et al., 2016; Buckman et al., 2015; Gao et al., 2017; Gao et al., 2014; Guillemot-Legris et al., 2016; Nadjar et al., 2017; Thaler and Schwartz, 2010; Thaler et al., 2012; Valdearcos et al., 2017; Valdearcos et al., 2014) we found that gene expression of the pro-inflammatory cytokine genes IL1β, IL6 and TNFα, as well as chemokine genes CCL2 and CCL5 was significantly increased in HFD-fed mice compared to SD-fed mice (Fig. 2A-E). The novelty of our study relies on the demonstration that an inflammatory-like gene response in the HT is physiologically induced by SD and arises within the first hours of food consumption. Moreover, this inflammatory-like gene response exhibits a diet specific profile defined by cytokine-/chemokine-type, amplitude and time course of the response, and is exacerbated in HFD-fed mice. In fact, the increase in IL-1β and TNFα gene expression appeared after 1h of food exposure (Fig. 2A and C) with a greater increase in HFD-fed mice for IL-1β (Fig. 2A). Moreover, the increase in CCL2 and CCL5 gene expression also appeared within 1h of food exposure, but was observed exclusively in HFD-fed mice (Fig. 2D and E). After 3h of food exposure, IL-1β, IL-6, TNFα and CCL2 gene expressions were increased only in HFD-fed mice (Fig. 2A-D). Finally, after 6h of food exposure, pro-inflammatory cytokine and chemokine gene expression in HFD-fed mice tend to return to SD-mice levels. Taken together, these results indicate that short term HFD exposure induces a specific and exacerbated hypothalamic inflammatory-like gene response, different from the one observed in SD-fed mice.

**Figure 2:**
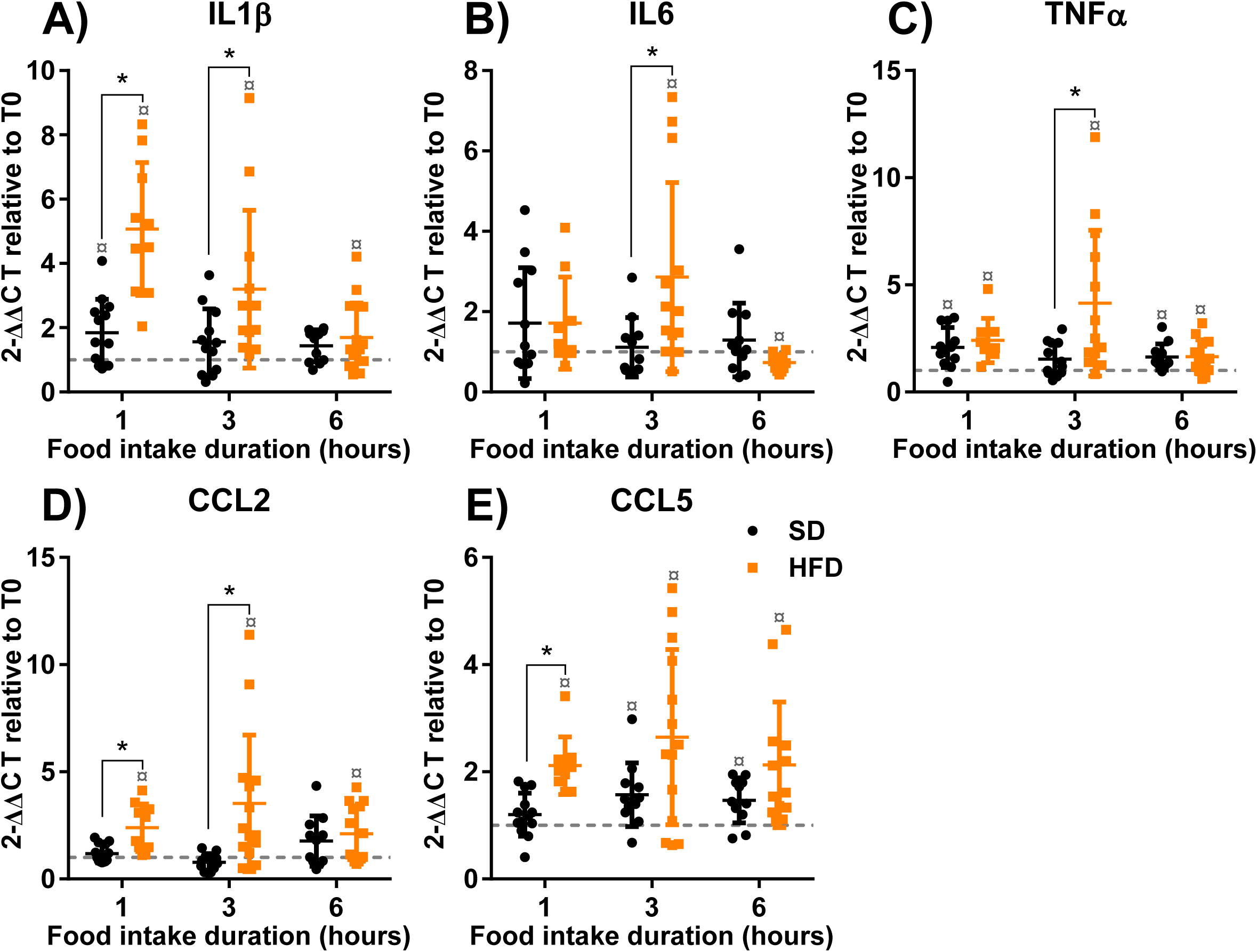
Short term high-fat diet exposure exacerbated post-prandial inflammatory-like gene response in the hypothalamus. Quantification of mRNA encoding proinflammatory cytokines IL-1β (A), IL-6 (B), TNFα (C) and chemokines CCL2 (D), CCL5 (E) in hypothalamus of 8-week-old C57Bl/6J male mice fed for 1h, 3h and 6h with either standard diet (SD) or high-fat diet (HFD). All mRNA levels were quantified relative to GAPDH housekeeping gene expression by the ΔΔCT method and presented as fold-change relative to baseline prior to diet exposure (T0, dashed line = initial level). *p < 0.05 Mann–Whitney test or unpaired t-test SD *vs* HFD; ¤p<0.05 Wilcoxon Signed Rank test SD/HFD *vs* 1; Mean+/- SD. N=9-12.

### 3.3. A single high-fat meal induces a specific acute neuropeptide gene expression profile in the hypothalamus

Given the acute and specific response in hypothalamic inflammatory-like gene expression of HFD-fed mice, we subsequently investigated the impact of a single high fat meal on hypothalamic neuropeptide genes. We examined changes in hypothalamic neuropeptide gene expression involved in energy balance regulation in the HT after 1h, 3h and 6h exposure to SD and HFD (Gao and Horvath, 2008). Food exposure induces changes in both SD- and HFD-fed mice. However, hypothalamic gene expression levels of orexigenic peptides MCH, AgRP and ORX (Fig. 3C, D and F) and anorexigenic peptide CART (Fig. 3E) were higher in HFD-fed mice compare to SD-fed mice while NPY levels remained equivalent in the 2 groups (Fig. 3A). In contrast, a decrease in anorexigenic peptide POMC gene expression, was observed at 1h only in HFD-fed mice (Fig. 3B). Interestingly, the time course of those changes was very specific for each diet. For example, SD-fed mice presented a transient and modest increase in MCH gene expression after 3h of food exposure compared to basal level (Fig. 3C) while it increased within the first hour of food exposure in HFD-fed mice and remained high for 6h (Fig. 3C). AgRP and ORX gene expression levels increased within the first hours of food exposure in both diet groups, with an exacerbated response in HFD-fed mice that lasted for 6h (Fig. 3D and F), unlike the increase in CART gene expression, which did not occur until 3h of food exposure and was only observed in HFD-fed mice (Fig. 3E). Taken together, our results show that short term HFD exposure is sufficient to specifically modulate hypothalamic neuropeptide gene expression involved in energy balance regulation, differently from what is observed in SD-fed mice.

**Figure 3:**
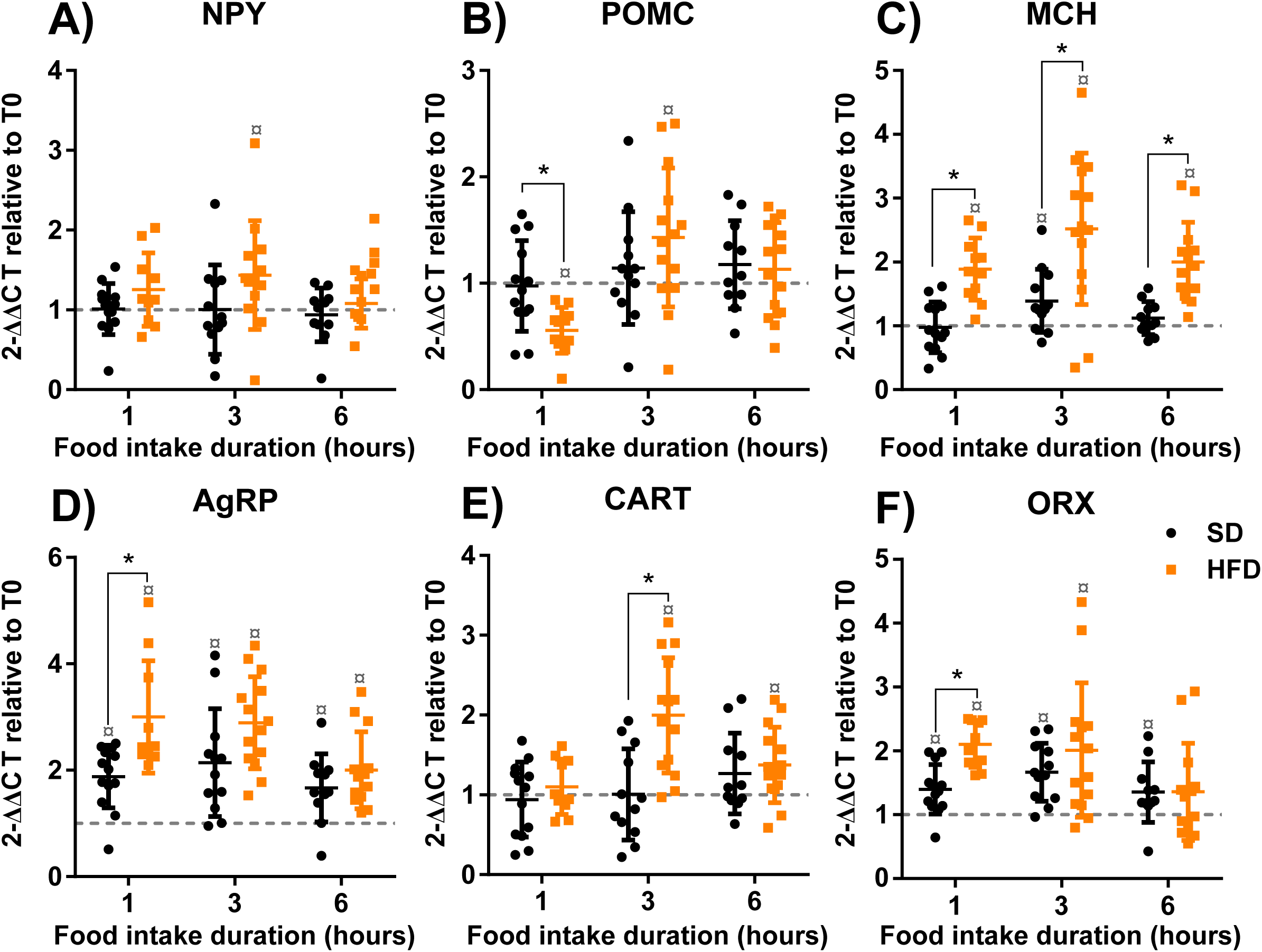
Short term high-fat diet exposure induces a specific acute neuropeptide gene expression profile in the hypothalamus. Quantification of mRNA encoding orexigenic neuropeptides NPY (A), AgRP (D), MCH (C), ORX (F) and anorexigenic neuropeptides POMC (B), CART (E) in hypothalamus of 8-week-old C57Bl/6J male mice fed for 1h, 3h and 6h with either standard diet (SD) or high-fat diet (HFD). All mRNA levels were quantified relative to GAPDH housekeeping gene expression by the ΔΔCT method and presented as fold change relative to baseline prior to diet exposure (T0, dashed line = initial level). *p < 0.05 Mann–Whitney test or unpaired t-test SD *vs* HFD; ¤p<0.05 Wilcoxon Signed Rank test SD/HFD *vs* 1; Mean+/-SD. N=9-12.

### 3.4. A single high-fat meal induces an acute increase of GFAP gene expression in the hypothalamus

Hypothalamic inflammation after medium-term (1 to 10 days) and long-term (2 to 32 weeks) HFD exposure is associated with astrogliosis (Balland and Cowley, 2017; Baufeld et al., 2016; Buckman et al., 2015; Thaler et al., 2012). It has yet to be determined whether the same is true at earlier time points. Given that our results showed an exacerbated hypothalamic inflammatory-like gene response in HFD-fed mice, we tested whether short-term exposure (within 1h, 3h and 6h) to HFD might affect astrocytes in the HT. To this end, we used the astrocyte marker GFAP for genetic and histological analysis. Surprisingly, 1h of food exposure increased GFAP gene expression in the HT of HFD-fed mice but not in SD-fed mice (Fig. 4A). Moreover, the increase in GFAP gene expression appeared to be transient as expression levels returned to baseline at 3h and 6h (Fig. 4A). We then tested if the increase in GFAP gene expression was associated with changes in astrocyte remodeling in the ARC, as described previously for medium term and long term HFD exposure (Balland and Cowley, 2017; Buckman et al., 2015; Thaler et al., 2012). SD and HFD-fed mice presented similar GFAP staining in the ARC, as shown in the photomicrographs displayed in Fig. 4C and corroborated by fluorescence quantification in the ARC of mice fed for 1h, 3h and 6h (Fig. 4B). These findings demonstrate that short term HFD exposure, specifically and transiently, upregulates hypothalamic GFAP gene expression.

**Figure 4:**
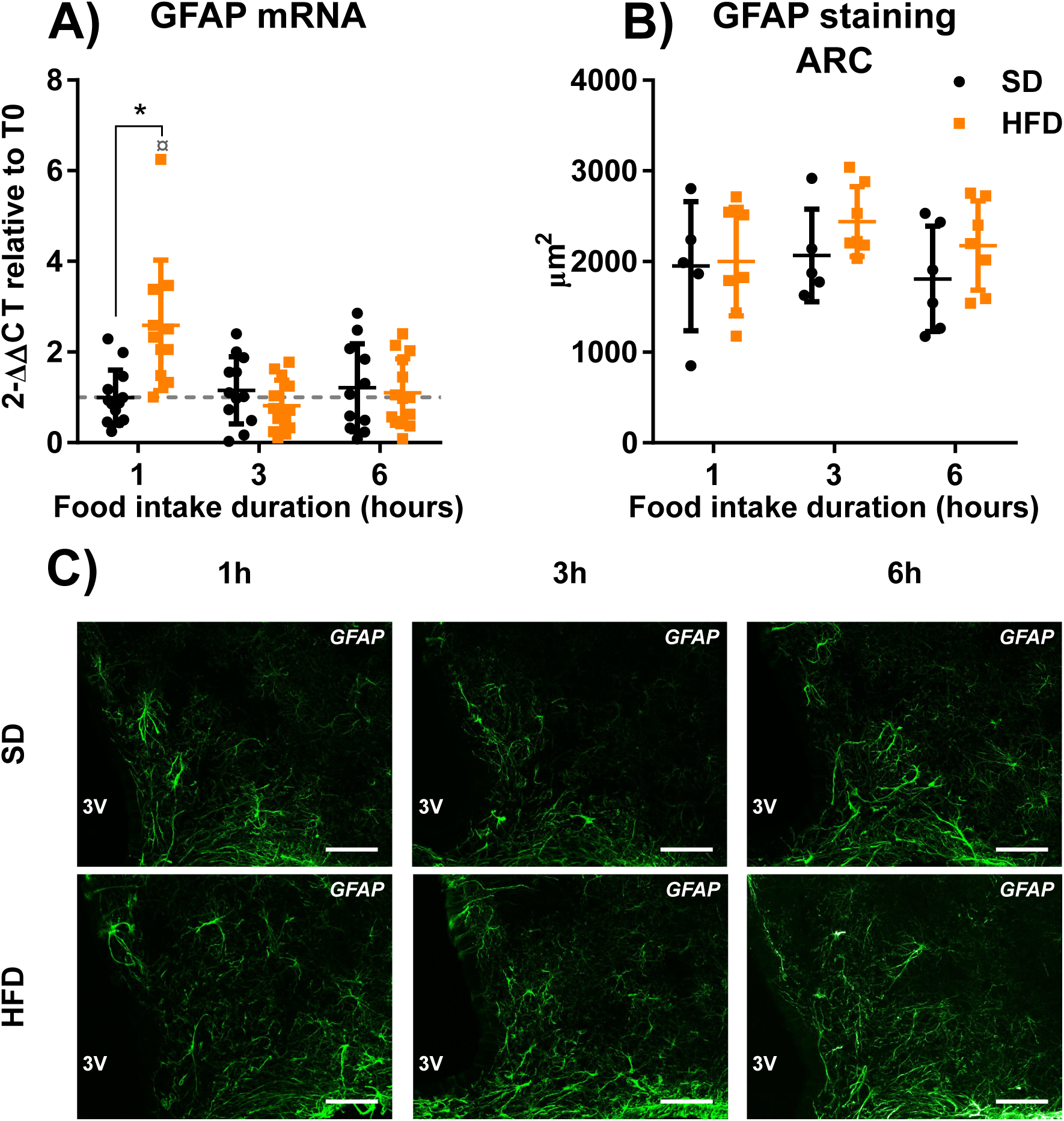
Short term high-fat diet exposure induces an acute increase of GFAP gene expression in the hypothalamus. Quantification of mRNA encoding the astrocyte marker GFAP (A) in hypothalamus of 8-week-old C57Bl/6J male mice fed for 1h, 3h and 6h with either standard diet (SD) or high-fat diet (HFD) N=9-12. Quantification of immunohistochemical detection of GFAP protein (stained area, μm^2^) (B) in coronal sections of the arcuate nucleus of hypothalamus (ARC; 30μm, −1.70mm relative to Bregma) from mice fed for 1h, 3h and 6h with either SD or HFD N=5-7. Immunohistochemical detection of GFAP protein in ARC of mice fed for 1h, 3h and 6h with either SD or HFD (C). All mRNA levels were quantified relative to GAPDH housekeeping gene expression by the ΔΔCT method and presented as fold-change relative to baseline prior to diet exposure (in A, T0, dashed line = initial level). Scale bar in (C): 50μm, 3V in (C): third ventricle *p < 0.05 Mann–Whitney test or unpaired t-test SD *vs* HFD; ¤p<0.05 Wilcoxon Signed Rank test SD/HFD *vs* 1; Mean+/- SD.

### 3.5. A single high-fat meal induces acute microglial remodeling in the arcuate nucleus of the hypothalamus

In addition to astrogliosis, hypothalamic inflammation after medium-term (3 to 7 days) and long-term (2 to 20 weeks) HFD exposure is also associated with microgliosis (Andre et al., 2017; Baufeld et al., 2016; Dorfman et al., 2017; Gao et al., 2017; Gao et al., 2014; Thaler et al., 2012; Valdearcos et al., 2017; Valdearcos et al., 2014). As for astrocytes, we tested whether short-term exposure (1h, 3h and 6h) to HFD might affect microglial cells in the HT. HFD exposure increased microglial marker Iba1 gene expression in the HT at 3h and 6h compared to SD-fed mice (Fig. 5A). We then tested whether Iba1 gene upregulation impacted Iba1 protein levels in the ARC, as previously described for medium term and long term HFD exposure (Andre et al., 2017; Baufeld et al., 2016; Dorfman et al., 2017; Gao et al., 2017; Gao et al., 2014; Thaler et al., 2012; Valdearcos et al., 2017; Valdearcos et al., 2014). No change in Iba1 immunofluorescence staining was observed (Fig. 5B and D). We also considered whether Iba1 gene expression modification might be associated with a change in microglial activity. Using the patch clamp technique, membrane properties of microglial cells were determined in 3h HFD-fed mice. We did not observe changes in the current-voltage relationship (Fig. 5E), but a significant increase in the cell capacitance in HFD-fed mice (Fig. 5F). We therefore looked more closely into cell morphology. While the histological approach may have had insufficient resolution to detect changes in soma size after 3h of HFD exposure (Fig. 5C and D), we did observe an increase in soma size in 6h HFD-fed mice (Fig. 5C), which is probably linked to the change observed in cell capacitance (Fig. 5F). Collectively, these results indicate that short term HFD exposure induces fast microglial cells remodeling in the ARC.

**Figure 5:**
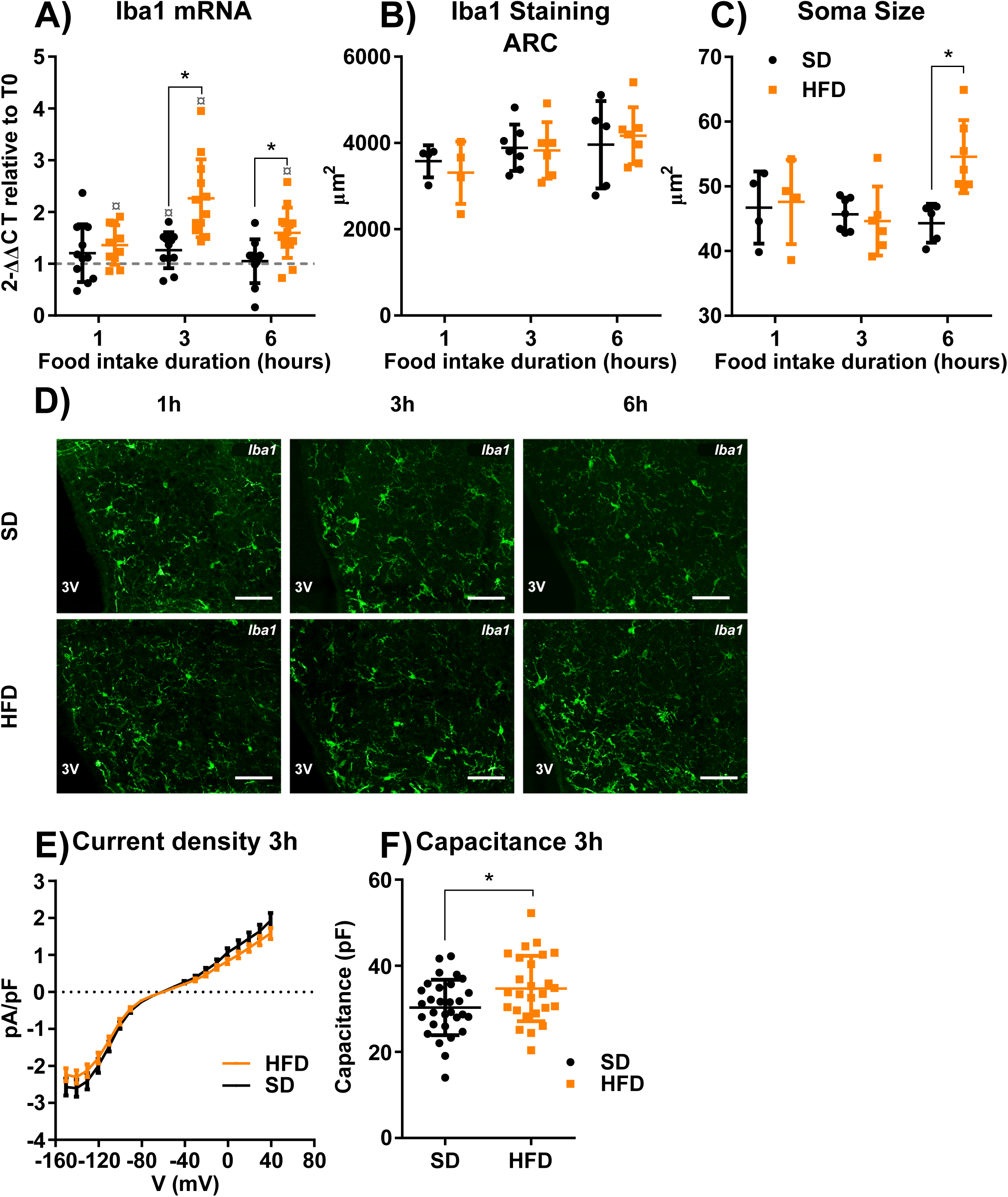
Short term high-fat diet exposure induces acute microglial remodeling in the arcuate nucleus of the hypothalamus. Quantification of mRNA encoding the microglial marker Iba1 (A) in the hypothalamus of 8-week-old C57Bl/6J male mice fed for 1h, 3h and 6h with either standard diet (SD) or high fat diet (HFD) N=9-12. Quantification of immunohistochemical detection of Iba1 protein (stained area, μm2) (B) in coronal sections of the arcuate nucleus of the hypothalamus (ARC) (30μm, −1.70mm relative to Bregma) from 8-week-old C57Bl/6J male mice fed for 1h, 3h and 6h with either SD or HFD N=4-7. Quantification of average microglial soma size (C) in ARC of 8-week-old C57Bl/6J male mice fed for 1h, 3h and 6h with either SD or HFD N=4-7 n=8-21. Immunohistochemical detection of Iba1 protein in ARC of 8-week-old C57Bl/6J male mice fed for 1h, 3h and 6h with either SD or HFD (D). Electrophysiological recording of current–voltage relationships (E) and membrane capacitance (F) of microglia cells whole-cell recorded in ARC of 9-10 week-old CX3CR1eGFP/eGFP male mice fed for 3h with either SD (N=3, n=30) or HFD (N=4 n=26). All mRNA levels were quantified relative to GAPDH housekeeping gene expression by the ΔΔCT method and presented as fold-change relative to baseline prior to diet exposure (in A, T0, dashed line = initial level). Scale bar in (D): 50μm, 3V in (D): third ventricle. *p < 0.05 Mann– Whitney test or unpaired t test SD *vs* HFD; ¤p<0.05 Wilcoxon Signed Rank test SD/HFD *vs* 1; Mean+/- SD.

### 3.6. Modulation of GFAP-positive cells changes post-prandial hypothalamic neuropeptide and inflammatory-like gene responses to SD and HFD exposure

We observed an acute and diet-specific hypothalamic gene response to food exposure for neuropeptides involved in energy balance regulation and pro-inflammatory markers. This response is associated with an increase in GFAP mRNA levels and microglial remodeling in the HT. We subsequently tested if modulating GFAP-positive cells and microglia could change this hypothalamic gene response. In order to modulate GFAP-positive cell activity we used DREADD (Designer Receptors Exclusively Activated by Designer Drugs) technology (Chen et al., 2016; Yang et al., 2015). An Adeno Associated Virus 8 (AAV8) with a GFAP promoter controlling DREADD Gi coupled with mCitrine gene sequence expression in the MBH of mice was used to target GFAP-positive cells. Analysis of mCitrine staining by immunohistochemistry confirmed successful injection (Supplementary Fig. 2A, B and C). Two weeks after virus injection, we treated mice with the specific agonist CNO (Clozapine N-Oxide) at a dose of 1 mg/kg to activate the DREADD Gi and 30 min later we exposed the mice to SD or HFD for 1h. Using this tool, we observed differential inflammatory-like and neuropeptide gene responses to SD and HFD in vehicle (NaCl) group (Fig. 6) compared to our original observation (Fig. 2 and 3). This could be explained by different inflammatory states of animals, as discussed later, but doesn’t change the originality of our results. CNO injection did not change 1h food consumption in both SD- and HFD-treated mice (Supplementary Fig. 2D and E). CNO injection did not alter POMC, MCH, AgRP and ORX gene expression response to SD and HFD exposure (Fig. 6B, C, D and F). In contrast, CNO injection, but not vehicle injection, decreased CART hypothalamic gene expression regardless of diet (Fig. 6E). Only NPY gene expression in response to food exposure is changed by the activation of DREADD Gi in GFAP-positive cells in a diet-specific manner (Fig. 6A). Interestingly, CNO injection, but not vehicle injection, decreased IL-1β, TNFα, and CCL2 gene expression in the HT regardless of diet (Fig. 6G, I and J). Moreover, CNO injection, but not vehicle injection, decreased CCL5 and Iba1 gene expression in HFD-treated mice (Fig. 6K and L). However, variability in CCL5 and Iba1 gene expression levels in SD-fed mice prevented statistically robust conclusions. CNO effect seems similar regardless of the diet on CCL5 gene expression (Fig. 6K), but the decrease in Iba1 gene expression is stronger in HFD-fed mice compared to SD-fed mice (Fig. 6L). Finally, IL6 gene expression in response to food exposure is changed by CNO injection in HFD-treated mice but not in SD-treated mice (Fig. 6H). Collectively, these results show that modulation of GFAP-positive cell activity in the MBH alters hypothalamic neuropeptide and inflammatory-like gene responses to both SD and HFD exposure. Interestingly, our results also demonstrate that specific NPY and IL-6 gene responses to food exposure are altered by the modulation of GFAP-positive cells activity in a diet-dependent manner.

**Figure 6:**
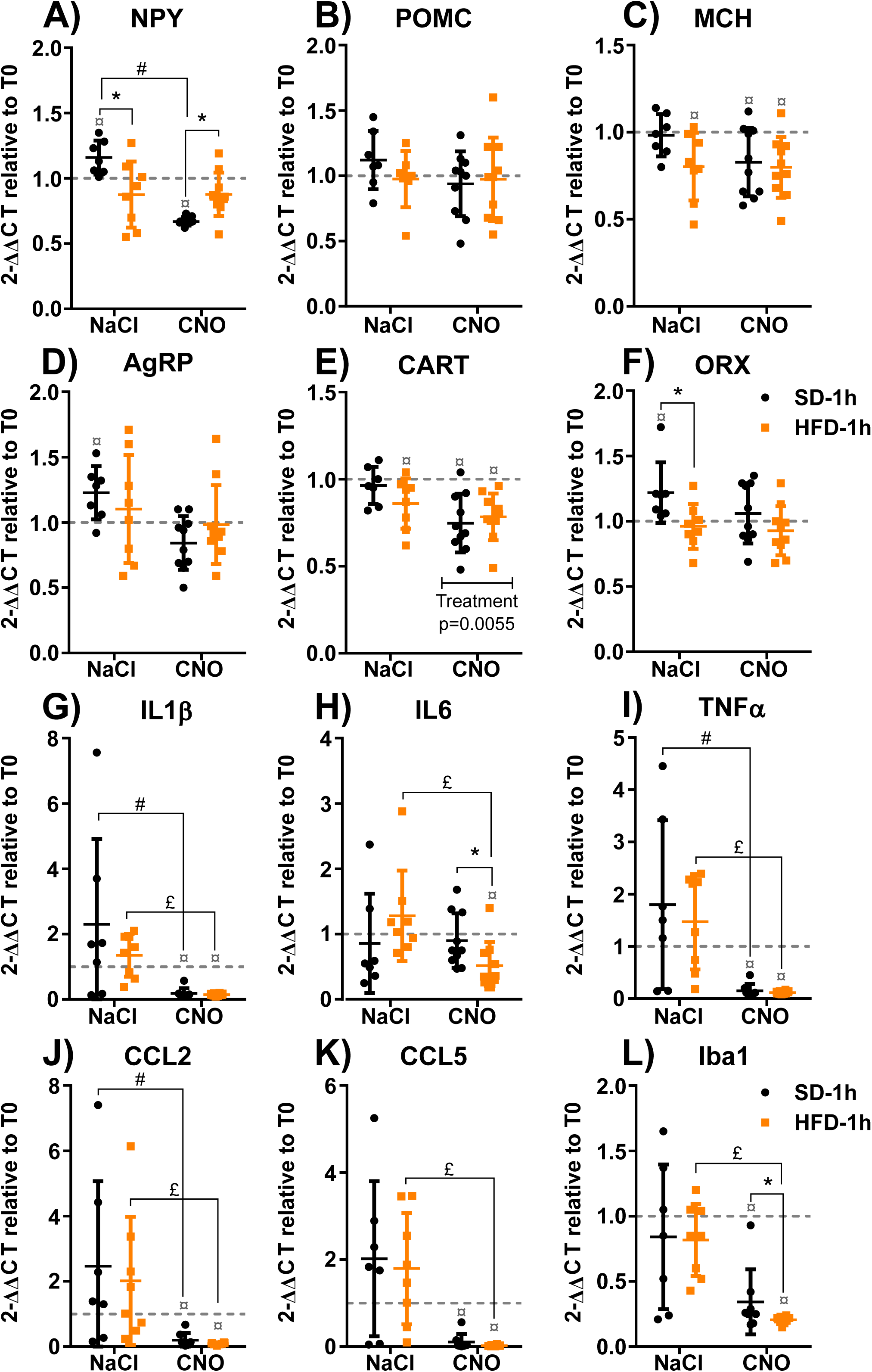
Modulation of GFAP-positive cells changes post-prandial hypothalamic neuropeptide and inflammatory-like gene responses to 1h SD and HFD exposure. Quantification of mRNA encoding orexigenic neuropeptides NPY (A), AgRP (D), MCH (C), ORX (F), anorexigenic neuropeptides POMC (B), CART (E), proinflammatory cytokines IL1β (G), IL6 (H), TNFα (I), chemokines CCL2 (J), CCL5 (K) and microglial marker Iba1 (L) in hypothalamus of mice fed for 1h with either standard diet (SD) or high-fat diet (HFD). 2 weeks prior to the experiment, 8-week-old C57Bl/6J male mice were injected with the AAV8-GFAP-DREADD-Gi virus in the mediobasal hypothalamus. To specifically modulate GFAP-positive cell activity, a selective DREADD-Gi receptor agonist, CNO (Clozapine-N-Oxyde), was injected intraperitonealy 30 min before food delivery (CNO). Control mice are injected with vehicle solution (NaCl). All mRNA levels were quantified relative to GAPDH housekeeping gene expression by the ΔΔCT method and presented as fold-change relative to NaCl injected-condition prior to diet exposure (T0, dashed line = initial level). *p ≤ 0.05 Mann–Whitney test or unpaired t-test SD *vs* HFD; ¤p ≤ 0.05 Wilcoxon Signed Rank test SD/HFD *vs* 1; #p ≤ 0.05 Mann–Whitney test or unpaired t-test NaCl-SD *vs* CNO-SD; £p ≤ 0.05 Mann–Whitney test or unpaired t-test NaCl-HFD *vs* CNO-HFD; Two-way ANOVA for multiple comparisons and interaction; Mean+/- SD. N=7-10.

### 3.7. Modulation of microglial activity changes post-prandial hypothalamic neuropeptide and inflammatory-like gene responses to SD and HFD exposure

To assess if microglia is involved in the neuropeptide and inflammatory gene responses induced by food exposure, we used PLX5622, which has been previously used to remove microglia from brain (Elmore et al., 2014). PLX5622 is a CSF1R agonist known to regulate microglial density in the brain. Two weeks after PLX5622 treatment, we assessed its efficiency to deplete microglia. PLX5622 almost completely ablated the microglial population as shown by the absence of Iba1 staining in the MBH (Supplementary Fig. 3A and B). We exposed PLX5622-treated and non-treated mice to SD or HFD for 1 h and 3h. Using this tool, we observed differential inflammatory-like and neuropeptide gene responses to SD and HFD in non-treated group (Fig. 7 and Supplementary Fig. 4) compared to our original observation (Fig. 2 and 3). This may be due to AIN76A diet exposure which might change inflammatory states as explained in the discussion, which does not undermine our results. SD and HFD intake (Supplementary Fig. 3E, F, H and I) and hypothalamic GFAP gene expression (Supplementary Fig. 3G and J) were not modified by PLX5622-induced microglial depletion. Moreover, PLX5622 treatment did not affect the hypothalamic neuropeptide gene response induced by 1h of food exposure (Supplementary Fig. 4A-F). PLX5622 treatment did not alter CART hypothalamic gene expression response to 3h food exposure neither (Fig. 7E). However, after 3h of food exposure, PLX5622 treatment increased similarly NPY and POMC gene expression regardless the diet (Fig. 7A and B). Concerning the AgRP gene response, result variability in SD-treated mice prevented a statistically robust conclusion, but PLX5622 treatment also seemed to upregulate AgRP gene expression regardless the diet (Fig. 7D). Interestingly, PLX5622 treatment changed MCH and ORX gene expression response to food exposure in a diet-dependent manner (Fig. 7C and F).

**Figure 7:**
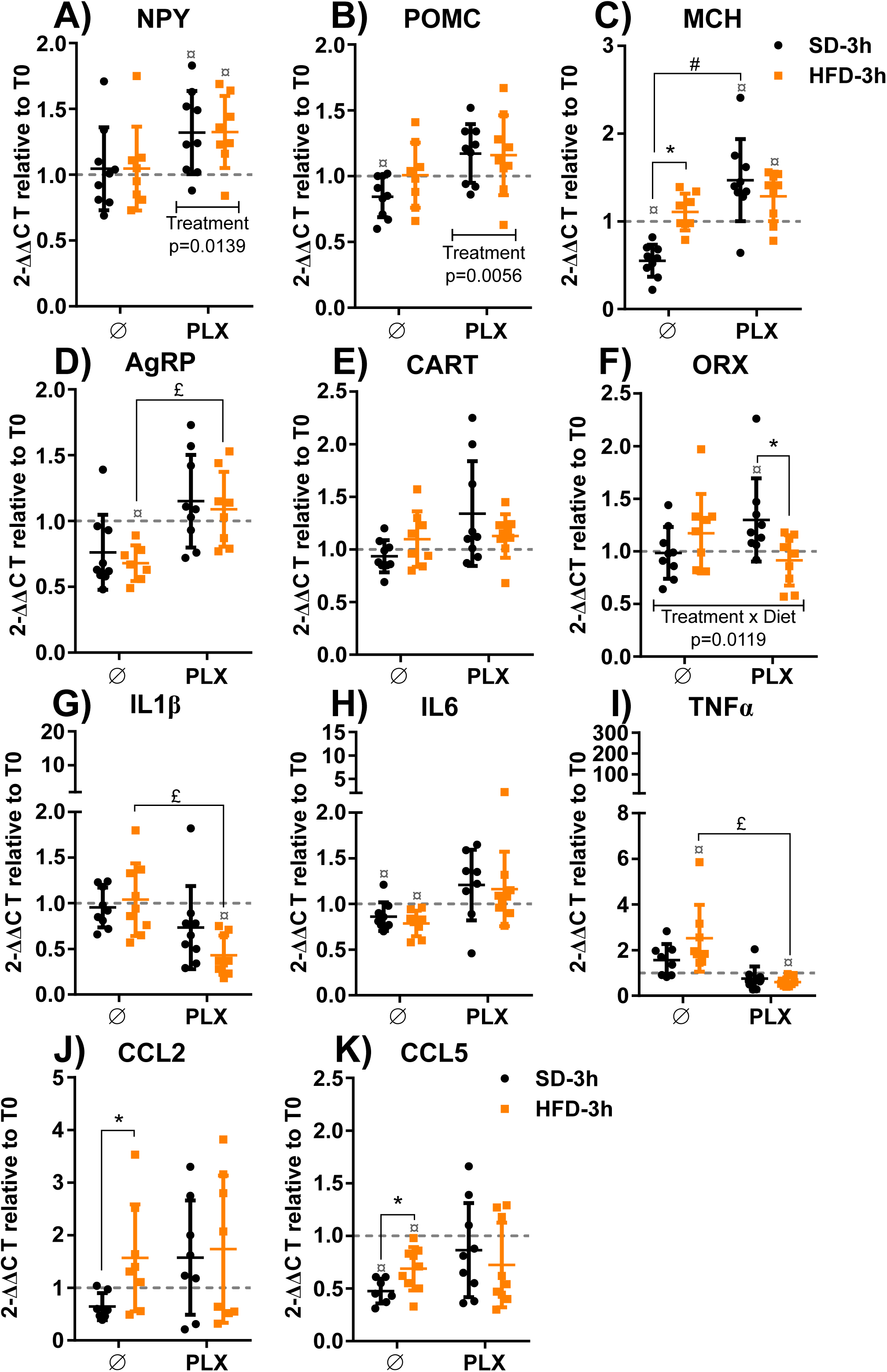
Modulation of microglial activity changes post-prandial hypothalamic neuropeptide and inflammatory-like gene responses to 3h SD and HFD exposure. Quantification of mRNA encoding orexigenic neuropeptides NPY (A), AgRP (D), MCH (C), ORX (F), anorexigenic neuropeptides POMC (B), CART (E), proinflammatory cytokines IL-1β (G), IL-6 (H), TNFα (I), chemokines CCL2 (J), CCL5 (K) in hypothalamus of mice fed for 3h with either standard diet (SD) or high-fat diet (HFD). Two weeks prior to the experiment, 8-week-old C57Bl/6J male mice were treated with the CSF1R (Colony Stimulating Factor1 Receptor) antagonist, PLX5622, formulated in the AIN76A food to remove microglia in the brain (PLX). Control mice received AIN76A food without PLX5622 for 2 weeks prior to the experiment (Ø). All mRNA levels were quantified relative to GAPDH housekeeping gene expression by the ΔΔCT method and presented as fold-change relative AIN76A-fed (+/-PLX5622) baseline prior to diet exposure (T0, dashed line = initial level). *p ≤ 0.05 Mann– Whitney test or unpaired t-test CHO *vs* HFD; ¤p ≤ 0.05 Wilcoxon Signed Rank test SD/HFD ≠ 1; #p ≤ 0.05 Mann–Whitney test or unpaired t-test Ø-SD *vs* PLX-SD; £p ≤ 0.05 Mann–Whitney test or unpaired t-test Ø-HFD *vs* PLX-HFD; Two-way ANOVA for multiple comparisons and interaction; Mean+/- SD N=8-9.

With respect to the impact of microglia depletion on inflammatory gene responses the high variance between the samples unfortunately detracted from any robust conclusion (Supplementary Fig. 4G-J and K; Fig. 7 G-K). Nevertheless, PLX5622 treatment increased TNFα mRNA levels in 1h HFD-treated mice (Supplementary Fig. 4I). Moreover, IL-1β and TNFα mRNA levels were decreased in 3h HFD-fed mice and effect seems similar on SD-fed mice (Fig. 7G and I).

Therefore, the results show that microglia depletion affects hypothalamic neuropeptide and inflammatory-like gene responses to both SD and HFD exposure. More interestingly, our results also demonstrate that specific MCH and ORX gene responses to food exposure are affected by the absence of microglial cells in a diet-dependent manner.

## 4. Discussion

Several studies have shown a link between HFD consumption, inflammation and risk factors associated with obesity. It seems certain that long-term exposure to HFD induces chronic low-grade systemic inflammation, which is responsible for the development of insulin resistance, type 2 diabetes mellitus along with other chronic illnesses such as cardiovascular disease or cancer (Cani et al., 2007; Emerson et al., 2017; Ghanim et al., 2009; Herieka and Erridge, 2014; Kelly et al., 2012; Lumeng and Saltiel, 2011; Thaler and Schwartz, 2010). More recently, hypothalamic inflammation, a process involving neurons, astrocytes and microglia, has been correlated with obesity and chronic HFD consumption (Balland and Cowley, 2017; Baufeld et al., 2016; Buckman et al., 2015; Gao et al., 2014; Guillemot-Legris et al., 2016; Kim et al., 2019; Milanova et al., 2019; Thaler and Schwartz, 2010; Thaler et al., 2012; Valdearcos et al., 2017; Valdearcos et al., 2014), underlining the effect of chronic exposure to HFD on the central inflammatory status. Interestingly, brain inflammatory responses in the HT were also observable after only one day of HFD consumption (Buckman et al., 2015) or 24h of lipid perfusion to the brain (Dalvi et al., 2017), well before the development of obesity. Moreover, inhibition of brain inflammatory responses to HFD leads to energy balance dysregulation (Buckman et al., 2015; Fernandez-Gayol et al., 2019). In humans, studies have described acute post-prandial inflammation associated to increased levels of circulating inflammatory markers in response to HFD (Emerson et al., 2017; Herieka and Erridge, 2014). Altogether, these data suggest that HFD-induced inflammation is a physiological response to cope with excess lipids, which could occur at the peripheral or central levels.

In our study, the calories from fat and sugar in HFD were equivalent (40% each), in contrast to numerous studies where fat content accounts for 60% kilocalories (kcal) while carbohydrate content was only 20% kcal. Therefore, the increase in caloric proportion represented by the lipids is more realistic in our model, and comparable to the typical human diet, where the occurrence of lipids in the presence of elevated carbohydrate content contributes to brain inflammation (Andre et al., 2017; Gao et al., 2017; Kim et al., 2019). According to previous studies of systemic post-prandial inflammation, HFD consumption promotes lipopolysacharide trafficking over the intestinal epithelium through trans-cellular and para-cellular transport (Valdearcos et al., 2014; Buckman et al., 2015). Analysis of endotoxin levels in the serum associated with inflammatory markers in both serum and HT, affirmed that post-prandial inflammation appears first in the HT. Therefore this study established, for the first time, the existence of a specific post-prandial hypothalamic inflammation response to both SD and HFD exposure linked to a unique modulation of mRNA levels of neuropeptides that are known to be involved in the regulation of energy balance (Gao and Horvath, 2008). However, while both SD and HFD consumption increased hypothalamic gene expression of the orexigenic neuropeptides AgRP and ORX, as well as inflammatory markers, IL-1β and TNFα, within the first hours of diet exposure, the amplitude of the response was higher in HFD-fed mice. Moreover, compared to SD exposure in the same time frame, 1h exposure to HFD induced selective expression of MCH orexigenic neuropeptide, as well as pro-inflammatory cytokines, CCL2 and CCL5. A sustained expression of proinflammatory cytokines IL-1β, IL-6, TNFα, chemokine CCL2 and orexigenic neuropeptide MCH was still observable after 3h exposure to food but only in HFD-fed mice. Finally, after 6h of food exposure, the differences observed between SD- and HFD-treated mice tended to disappear, except for the MCH orexigenic neuropeptide mRNA level, which remained higher in the HFD group. This suggests that short-term exposure to a lipid-rich diet triggers hypothalamic gene expression of orexigenic neuropeptides leading to an adaptive behavioral response correlated to increased kilocalorie intake observed 6h post HFD exposure (Gao and Horvath, 2008). It is not yet known how these genetic changes occur in orexigenic neurons, although we believe that they may be the consequence of lipid-induced neuroinflammation. We have yet to define the role of the inflammatory brain cells in this response over such a short time frame in this study. Nevertheless, a recent study demonstrated that one day of lipid perfusion to the brain is sufficient to activate an inflammatory response in NPY/AgRP neurons through TNFα and ER-stress signaling (Dalvi et al., 2017). We could therefore expect such a mechanism to appear even earlier in response to HFD, which could modulate the neuropeptide expression involved in energy balance regulation as we observed in our study.

As previously described, HFD-induced hypothalamic inflammation is associated with glial reactivity (Balland and Cowley, 2017; Baufeld et al., 2016; Buckman et al., 2015; Gao et al., 2014; Guillemot-Legris et al., 2016; Kim et al., 2019; Milanova et al., 2019; Thaler et al., 2012; Valdearcos et al., 2017; Valdearcos et al., 2014; Waise et al., 2015). First, we focused on GFAP-positive cells. Targeting of cell activity decreases the postprandial pro-inflammatory cytokines IL-1β, IL-6, TNFα and the chemokines CCL2 and CCL5 gene response to both SD and HFD exposure. Interestingly AAV8-DREADD-Gi-mCitrine injection exclusively in MBH is powerful enough to downregulate pro-inflammatory marker mRNA levels measured in the whole HT, suggesting that MBH GFAP-positive cells play a critical role in hypothalamic post-prandial inflammation. Moreover, we have provided novel evidence that an acute exposure to HFD specifically increases GFAP mRNA levels in HT within the first hours of food consumption. Another observation worth highlighting is that the IL-6 gene response to food exposure is modulated by GFAP-positive cells in a differential way, depending on diet. In fact, modulation of GFAP-positive cellular activity decreases IL-6 mRNA hypothalamic levels only in HFD-treated mice. This is consistent with recent discoveries demonstrating that IL-6 produced by astrocytes is a major actor in central energy balance regulation and plays a protective role against obesity development during HFD exposure (Fernandez-Gayol et al., 2019; Quintana et al., 2013; Timper et al., 2017). On the other hand, modulation of GFAP-positive cellular activity does not dramatically affect hypothalamic neuropeptide mRNA levels and does not change HFD intake. This is consistent with previously published results and indicates that modulation of NPY/AgRP neuron activity by GFAP-positive cells in response to HFD occurs over a different time frame from our study (Chen et al., 2016; Yang et al., 2015).

Because glial reactivity to HFD consumption is also characterized by microglial reactivity, we then focused on microglia. In accordance with previous studies, we did not observe any effect on HFD intake when microglia was depleted (Waise et al., 2015). We found that 3h of HFD exposure increases hypothalamic mRNA levels of the microglial marker Iba1 compare to SD-fed mice. This rise was associated with soma size enlargement of microglial cells in the ARC of HT only in HFD treated mice. For microglial cells, the association of a specific activity state with morphological aspects remains controversial (Kettenmann et al., 2011; Tremblay et al., 2011). However, what is certain is that soma size enlargement of microglial cells and the modification of Iba1 gene expression initiated by HFD consumption reflects a change in their activity, which might enable protective mechanisms to deal with lipid excess (Kim et al., 2019; Thaler et al., 2012; Valdearcos et al., 2017; Valdearcos et al., 2014). Here we provide novel evidence that microglial reactivity to HFD exposure appears within the first hours of food exposure. Moreover, our approach clearly shows that microglial cells participate in the post-prandial hypothalamic inflammation induced by both SD and HFD, though it is too early to state to what extent and what diet-specificity. In fact, microglial depletion seems to exacerbate hypothalamic inflammatory-like response to 1h HFD exposure, but that amplitude varies between individuals. Because the variability cannot be explained with a correlation with body weight and food intake (data not shown) nor by PLX efficiency (Supplementary Fig. 3) it is tempting to speculate that the involvement of microglia in HFD-induced hypothalamic inflammation differs between subjects and may be involved in the predisposition of an individual to the HFD-induced hypothalamic disturbances involved in obesity. Moreover, our experiments showed that microglial depletion modulated gene response to food exposure of neuropeptides involved in energy balance regulation. Not only did it increase NPY and POMC hypothalamic mRNA levels in both SD- and HFD-treated mice, but it also differentially affected MCH and ORX gene responses to diet type, suggesting that microglial responses to food exposure could specifically act on neurons involved in energy balance regulation in a diet-dependent manner.

Up to now, astrocytic or microglial reactive gliosis in the ARC of the HT in response to chronic HFD exposure and their participation in HFD-induced inflammatory processes and functional impact is still controversial and may depend on the nutritional lipid composition (Harrison et al., 2019; Kim et al., 2019; McLean et al., 2019; Valdearcos et al., 2014). Our study offers new insight on the impact of HFD consumption on astrocytic or microglial populations located in areas involved in energy balance regulation. Both cell types may be activated by short-term food exposure and orchestrate inflammatory-like processes in the ARC modulating neuronal function, but in differential time frames and with a diet specificity. Indeed, during short-term HFD exposure, GFAP gene expression is acute and transient, whereas Iba1 gene expression and microglial reactivity occurs after 3h of diet exposure and lasts over 6h. Moreover, the time frame of microglial reactivity to short-term HFD exposure coincides with the appearance of endotoxin and saturated fatty acid-enriched triglyceride serum content. This supports the hypothesis that saturated fatty acid-induced hypothalamic inflammation may require microglial activation (Kim et al., 2019; Valdearcos et al., 2014). Therefore, our experiments suggest that the increase of lipid content in diet is primarily sensed by GFAP-positive cells, which initiates the hypothalamic inflammatory response followed by hypothalamic microglial cell activation by central and systemic signals that impact energy balance regulation through the modulation of hypothalamic neuropeptide expression.

Despite the originality of our results, we are aware of study limitations. While we focused on the HT because it is a well-known area involved in energy balance regulation (Gao and Horvath 2008), we do not exclude the possibility that other brain areas may display post-prandial inflammation associated with neuropeptide modulation. Indeed, several studies have shown that chronic HFD consumption leads to hippocampal inflammatory responses and affects behavioral disorders associated with obesity (Waise, Toshinai et al. 2015; Guillemot-Legris, Masquelier et al. 2016; Gzielo, Kielbinski et al. 2017; Tsai, Wu et al. 2018; Spencer, Basri et al. 2019). Furthermore, our methodology prevented a detailed analysis of subtle changes that might occur in the several functionally distinct nuclei of the HT. Moreover, to gain insight on the involvement of each cell type, we used two different tools to manipulate GFAP-positive and microglial cell activity which must be called into question. DREADD technology has already been used to target GFAP-positive cells of the ARC glia in the regulation of neuronal subtype-specific modulation of energy homeostasis (Yang, Qi et al. 2015; Chen, Sugihara et al. 2016). Given this context, we initiated our study using DREADD Gi as this strategy was proven to achieve cellular inhibition - at least in neuronal cell types (Roth 2016). However, a 2019 study has demonstrated that Gi/o protein-coupled receptor may differentially affect brain cell type, inhibiting neurons and activating astrocytes stimulating gliotransmitter release (Durkee, Covelo et al. 2019). Thus, at this stage while it seems difficult to definitely conclude whether GFAP-positive glial cell activation or inhibition mediates HFD-induced feeding behavior changes, our results clearly support GFAP-positive glial cell involvement in this process. Concerning microglial cells, although new tools using DREADD technology to target microglia have been recently reported (Grace, Wang et al. 2018), we chose PLX5622 as it was the only tool available at the time we initiated the study to modulate microglial cells in the brain. PLX5622 has been shown to act mainly on microglia in adult whole brains (Elmore, Najafi et al. 2014; Oosterhof, Kuil et al. 2018). Thus, in our study we analyzed total brain microglial depletion effects, and recognize that further analysis needs to be done to specify microglial involvement in hypothalamic post-prandial inflammation. Besides, using those two tools we observed differential inflammatory and neuropeptide gene responses in respective control groups for AAV8-DREADD-Gi-mCitrine or AIN76A diet experiments compared to our original observation without cellular targeting. This could be explained by different inflammatory states of animals: sham and non-treated animals vs injected with virus vs treated with an AIN76A diet containing PLX5622. While it is obvious that stereotaxic injection of virus has immunological implications that could affect brain inflammatory status, it is worth noting that 76.9% of carbohydrates contained in AIN76A are sucrose, known to play a critical role in diet-induced hypothalamic inflammation (Andre, Guzman-Quevedo et al. 2017; Gao, Bielohuby et al. 2017). The WT mouse, the mice injected with the virus and the mice exposed to AIN76A can therefore be considered as three different study models in which we analyzed hypothalamic post-prandial inflammation and glial cells involvement in this process.

Overall, while the effect of medium-term and long-term nutritional lipid exposure has been extensively demonstrated to promote hypothalamic inflammation involving astrocytic and microglial cells (Andre et al., 2017; Balland and Cowley, 2017; Baufeld et al., 2016; Gao et al., 2017; Gao et al., 2014; Guillemot-Legris et al., 2016; Nadjar et al., 2017; Thaler et al., 2012; Valdearcos et al., 2017; Valdearcos et al., 2014), little was known about short-term exposure. Here we demonstrate, for the first time, that a moderate increase of nutritional lipid content in diet – representative of current human diets – can exacerbate hypothalamic post-prandial inflammation in a short time and that this response involves hypothalamic GFAP-positive cells and microglial cells. Although this primary response is probably aimed at regulating energy balance, continual nutritional lipid excess may contribute to dysregulating those signals and contribute to the hypothalamic disturbances leading to obesity (Thaler and Schwartz, 2010). Because the amplitude of systemic post-prandial inflammation has been correlated with type 2 diabetes and cardiovascular diseases (Emerson et al., 2017; Milan et al., 2017), our study leads us to believe that exacerbated hypothalamic post-prandial inflammation might predispose individuals to obesity and needs to be characterized in order to address this worldwide crisis.

## Acknowledgments

We thank Plexxikon and Dr. Brain West for PLX5622. This work has been supported by the CNRS, the Fondation pour la Recherche Médicale (DEQ20150331738), the Medisite Foundation and by the French government, managed by the National Research Agency (ANR), through the UCA^JEDI^ Investments in the Future project with the reference number ANR-15-IDEX-01, the LABEX SIGNALIFE program with the reference number ANR-11-LABX-0028-01 and La Fondation Nestlé France. We thank Dr. JP. Pais de Barros from the Lipidomic Platform uBourgogne (INSERM UMR1231 / LabEX LipSTIC) for 3OH C14:0 blood analysis. We thank Abby Cuttriss (Office of International Scientific Visibility, Université Côte d’Azur), Dr. Merel Rijnsburger (VU University Medical Center Amsterdam) and Pr. Christophe Magnan (Paris Diderot University) for editing the manuscript.

## Author contributions

C.C., J.L.N., N.B. and C.R. conceived and supervised the study, designed experiments, and wrote the manuscript. CC. performed the majority of the experiments, interpreted results, and generated figures. K.S. performed EIA and multiplex assays. O.L.T. initiated immunohistochemistry experiments and quantitative PCR experiments with CL. J.L. performed most of the quantitative PCR experiments and contributed to data analysis. C.A.M. performed and analyzed results from electrophysiology experiments. S.BF. performed food intake experiments. N.D. participated to management of mice lines. D.D and L. F. performed TG measurements. F.B. participated to the quantification of astrocytes and microglia morphology changes. E.A. and A.B. contributed to data analysis and paper writing. All authors discussed results and/or reviewed the manuscript.

## Conflict of interest

The authors declare no conflict of interest.

## Abbreviations

ACN: Acetonitrile
AgRP: Agouti-Related Peptide
ARC: Arcuate nucleus
CART: Cocaine- and Amphetamine-Regulated Transcript
CCL: Chemokine (C-C motif) ligand
ChCl3: Chloroform
CNO: N-Oxide Clozapine
CSF1R: Colony Stimulating Factor 1 Receptor
DREADD: Designer Receptors Exclusively Activated by Designer Drugs
GAPDH: GlycerAldehyde Phosphate DeHydrogenase
GFAP: Glial Fibrillary Acidic Protein
HFD: High-Fat Diet
HT: Hypothalamus
Iba1: Ionized Calcium Binding Adaptor Molecule 1
IL: Interleukin
INF: Interferon
IPA: Isopropanol
MBH: Medial Basal Hypothalamus
MCH: Melanin-Concentrating Hormone
MeOH: Methanol
NPY: NeuropeptideY
ORX: Orexine
PBS: Phosphate Buffer Saline
POMC: Pro-OpioMelanocortin
SD: Standard Diet
TG: Triglyceride
TNF: Tumor Necrosis Factor.

## Supplementary data

Supplementary data associated with this article can be found in this online version.

